# Extracellular vesicles from monocyte/platelet aggregates modulate human atherosclerotic plaque responses

**DOI:** 10.1101/841361

**Authors:** S Oggero, M de Gaetano, S Marcone, M Barry, T Montero-Melendez, D. Cooper, L V Norling, E P Brennan, G Godson, M Perretti

## Abstract

In atherosclerosis, a chronic disease characterized by lipid accumulation, fibrosis and vascular inflammation, extracellular vesicles (EVs) are emerging as key players in different stages of disease development. Here we provide evidence that EVs released by mixed aggregates of monocytes and platelets in response to TNF-α are both CD14+ and CD41+. Tempering platelet activation with Iloprost™ impacted the quality and quantity of EV produced. Proteomics of EVs from cells activated with TNF-α alone or in presence Iloprost™ revealed distinct proteome, with selective hits like gelsolin. EVs from TNF-α stimulated monocytes augmented release of cytokines, and modulated more than 500 proteins by proteomics, when added to human atherosclerotic plaques. In contrast, EVs generated by TNF-α and Iloprost™ produced minimal plaque activation. In conclusion, attenuating platelet activation has an effect on EV composition released from monocyte/platelet aggregates with downstream modulation of their pro-inflammatory actions and contribution to the development and progression of atherosclerosis.

## Introduction

Extracellular vesicles (EVs) are cell-borne particles that contain a complex biological cargo composed of nucleic acids, proteins and lipids. Firstly described by Wolf in 1967, EVs were reported to have prothrombotic functions (1) and proposed to be a way for cells to dispose of unnecessary products (2). Since then, EVs have been ascribed extended properties impacting on both pathological and physiological processes, including modulation of adaptive immune response (3), tumour metastasis and growth (4), and coagulation cascade (5).

The majority of work conducted so far with EVs has focused on identifying the markers of cell of origin they bear; however, using neutrophil-derived EVs as prototypes, we proposed that their composition and hence properties would vary to reflect the environment surrounding the cell source (6). This study helped to develop the concept of EV heterogeneity (7): EVs mirror the activation state of their cell of origin through specific enrichment or presence of given proteins, lipids and nucleic acids, affecting in this manner their biological properties. For example, the procoagulant property of endothelial cell-derived EV is largely dependent on both the exposure of tissue factor and phosphatidylserine on the particle surface (8, 9), while EV-mediated induction of endothelial cell proliferation is mainly tissue factor-dependent (10).

Atherosclerosis is the most prominent and common cause of cardiovascular diseases responsible for ∼50% of all deaths in Europe (11). Complications of atherosclerosis, especially acute coronary syndromes, have been linked to rupture of vulnerable lesions, causing atherothrombosis and vessel occlusion. In the pathogenesis of atherosclerosis, most of the cellular and molecular events including endothelial dysfunction, platelets activation and monocyte and macrophage accumulation, have been characterized (12), yet effective prevention of atherosclerosis and adverse cardiovascular events are still needed. Thus, studying the lesion biology is essential for growing our knowledge on the pathophysiology of atherosclerosis and to allow identification and development of novel therapeutic strategies. In this context, the possible implication of EVs in promoting and progressing this pathology is a recently explored field.

There is evidence for EVs to cause endothelial dysfunction, vascular calcification, unstable plaque progression, rupture and thrombus formation (13). Regarding plaque formation and destabilization, studies have focused on plaque-released EVs. For example, atherosclerotic plaque EVs expressed surface antigens of leukocyte origin (including major histocompatibility complex classes I and II), and promoted T-cell proliferation (14). In terms of EVs effects once added to the plaque, there is *in vivo* evidence for monocyte EVs to promote leucocyte adhesion to post-capillary venules and T-cell infiltration in atherosclerotic plaques (15). The majority of these studies have been conducted with murine models and *in vitro* cellular assays. However a better assessment of the inflammatory processes in human atherosclerosis can be attained through organ culture approaches, rather than using less complex experimental settings.

Here, we characterise human monocyte-derived EVs particularly in presence of platelets, to mimic a vascular inflammatory status, and define the composition of these EVs and their biological function once added to human atherosclerotic plaques, observing a positive feed-forward mechanism fuelling inflammation and possibly instability. Intriguingly, attenuating platelet activation has an impact on EV composition and a functional effect on modulating the reactivity of the atherosclerotic plaque.

## Results

### Monocyte-derived EVs are regulated by aggregation with platelets

After preparation of an enriched population of monocytes from human blood using negative selection procedure, flow cytometry analysis demonstrated a high degree of monocyte/platelet aggregates whereby 56.5±5.1% of CD14+ events were also positive for CD41+, the platelet marker (n=10; Fig. 1a and b). In order to determine whether platelet-monocyte interactions were dependent on platelets activation, we introduced Iloprost a potent analogue of prostacyclin in our isolation protocol. Addition of Iloprost during the isolation procedure did not impact on the formation of these aggregates (Fig. 1c), a phenomenon also visualised by ImageStream™ (Fig. 1f). A similar outcome was observed when cells were purified using the Histopaque low density gradient protocol (Fig. 1c).

**Figure 1.**
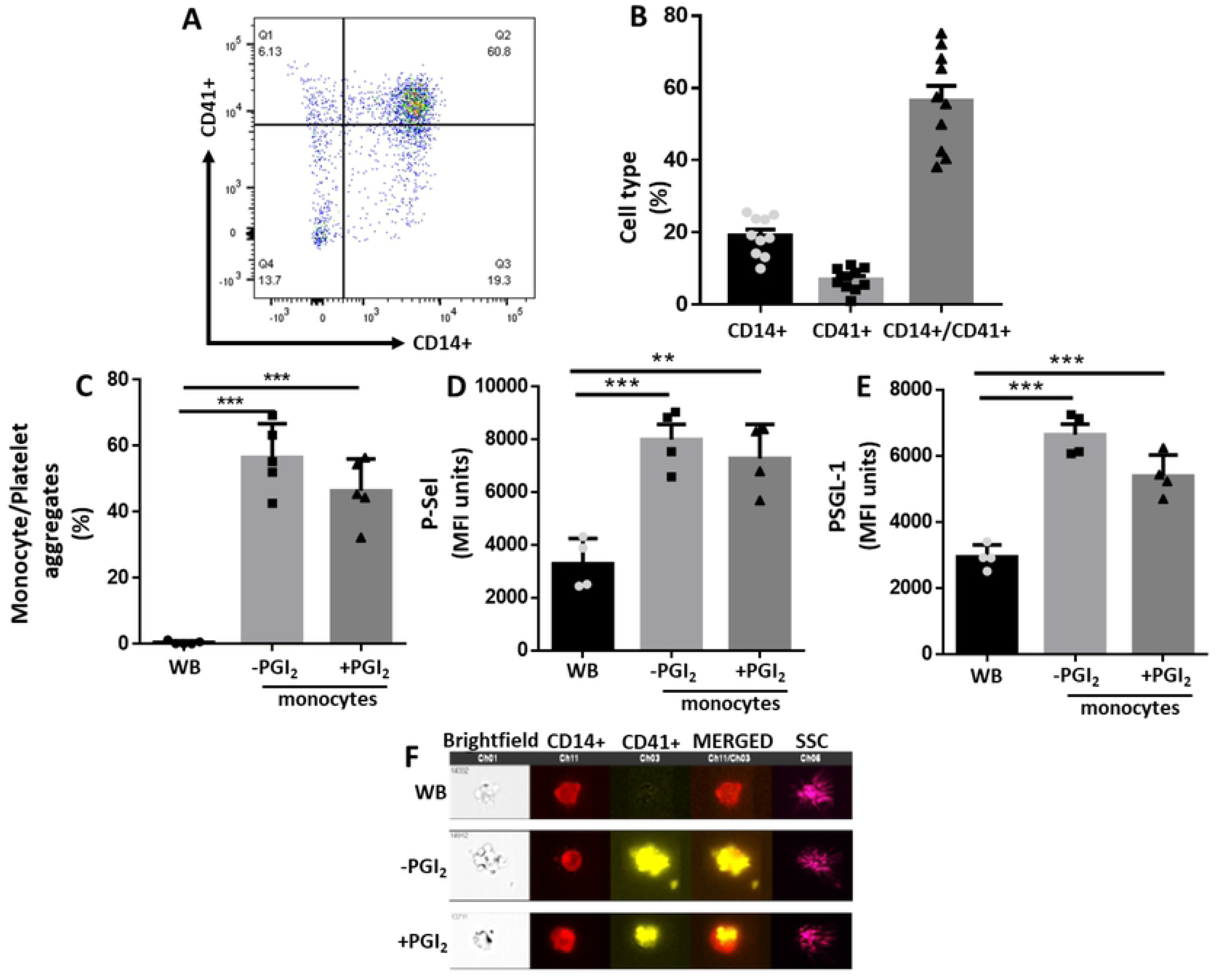
Iloprost controls platelet but not monocyte activation. Whole blood (WB) or monocytes isolated using the RosetteSep purification protocol were incubated with or without 1 µM Iloprost (PGI_2_). CD14 and CD41 were used as markers for monocytes and platelets, respectively, for flow cytometry and Imagestream™ analyses. (a) Dotplot graph showing monocyte (CD14+) and platelet (CD41+) immunostaining to reveal presence of aggregates (double positive events). (b) Percentages of CD14+ (monocytes), CD41+ (platelets), CD14+/CD41+ double positive (aggregates) events. Data are mean ± SEM of n=10 distinct cell preparations. Representative of n=10 distinct cell preparations. (c) Proportion of aggregates as analysed by Imagestream™ comparing Whole blood (WB) cells with purified monocytes in the presence or absence of Iloprost (*p<0.05, **p<0.01, ***p< 0.001; one-way ANOVA post Bonferroni test, mean ± SEM, n=5 distinct preparations). (d) P-selectin expression of monocyte/platelets aggregates following presence or absence of Iloprost compared to WB cells. Data are mean ± SEM of n=4 distinct cell preparations. (e) PSLG-1 expression of monocyte/platelet aggregates upon addition or not of Iloprost following presence or absence of Iloprost compared to WB cells. Data are mean ± SEM of n=4 distinct cell preparations. (f) Visualization of monocyte/platelet aggregates as identified by ImageStream™. Representative of n=5 distinct cell preparations.

A degree of monocyte and platelet activation consequent to the purification procedure was confirmed by transient cell surface expression of P-selectin and PSGL-1 compared when cells were purified using the Histopaque low density gradient protocol (Fig. 1d and e). EV-free monocyte supernatants were analysed to assess activation status, and an increase in all cytokine and chemokine levels were detected with TNF-α treatment with no significant modulation by the prostacyclin analogue (Fig S1).

These data indicate that monocyte isolation leads to immune cells carrying platelets and that Iloprost addition does not affect the extent of this interaction. Since monocyte/platelet aggregates are typical of several cardiovascular settings, including atherosclerosis (see Discussion), we decided to exploit this enriched monocyte preparation herein obtained to study formation and properties of EVs generated in these cell-to-cell crosstalk settings.

For deep analysis of EVs, we implemented a validated protocol where fluorescence triggering of EVs (labelled with BODIPY-FITC) allows a better identification by ImageStream™(Headland et al. 2015). Using a double gating strategy for staining with CD14+ and CD41+, EVs from platelets (CD41+/CD14-; ∼15%), monocytes (CD41-/CD14+; ∼60%) and a subset bearing both markers (CD41+/CD14+; ∼7.5%) were identified, both in presence and absence of TNF-α and Iloprost (Fig. 2a). TNF-α addition to monocytes almost doubled the number of total EVs compared with unstimulated cells (n=5, P<0.01) (Fig. 2b). Addition of Iloprost did not affect basal EV numbers (Fig. 2b). Similar results were obtained for total CD14+ EV, however addition of Iloprost significantly reduced (∼40%) the proportion and number of both CD41+/CD14- and CD14+/CD41+ EVs as quantified in response to TNF-α stimulation (Fig. 2c-e). When the cellular preparations were stimulated with PAF, a known activator of platelets as well as of monocytes, a larger number of EVs were produced with a higher proportion of CD14+/CD41+ events (1/5 of total CD14+ events); this time Iloprost afforded a marked reduction of all EV subset values (Fig. S2). Thus while Iloprost did not affect formation of monocyte/platelet aggregates and exerted selective inhibition on TNF-α stimulated EV numbers and phenotypes, it markedly affected PAF stimulation indicating high efficacy in reducing platelet and monocyte activation, regulating mainly the release of platelet EVs (CD41+) and CD14+/D41+ double positive EVs.

**Figure 2.**
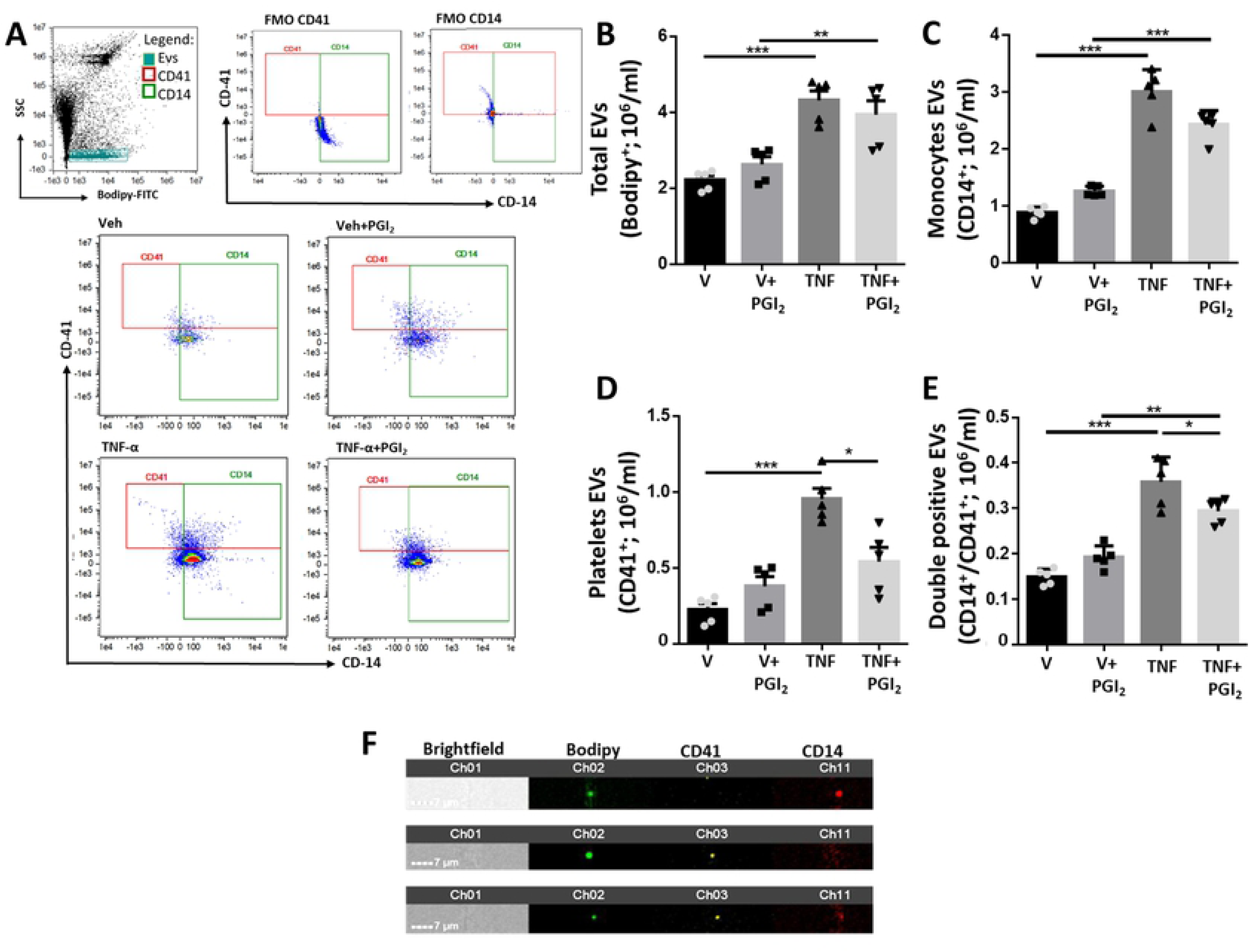
Characterization of monocyte/platelet derived EVs. Monocytes were isolated using the RosetteSep purification protocol. Cells (1×10^6^/ml) were incubated with vehicle (V) or TNF-α (50 ng/ml), in presence or absence of Iloprost (1 µM; PGI_2_) for 60 min. (a) ImageStream™ analysis of the vesicle showing quadrant selections, FMOs and representative images following staining for anti-CD14 or anti-CD41. EV generation in cell-free supernatants was quantified following Bodipy staining for total vesicles (b); monocyte CD14+ EVs (c); platelet CD41+ EVs (d) and double positive CD14+/CD41+ vesicles (e). (*p<0.05, **p<0.01, ***p< 0.001; one-way ANOVA post Bonferroni test, mean ± SEM, n=5 distinct preparations). (f) Visualization of CD14+ (top panel), CD41+ (middle panel) and CD14/CD41 double positive (bottom panel) EVs by ImageStream™.

The physical characteristics of EVs were studied by nanoparticle tracking analysis. This set of experiments demonstrated that vesicles produced in these settings ranged between 50  and 500 nm in diameter (Fig. S3); addition of Iloprost had modest or nihil effect on the physical characteristics of the EV samples. All preparations of EVs displayed similar size mean and mode regardless of the stimulating agent applied or presence of Iloprost (Fig. S3).

### EVs differentially activate HUVEC

Since EVs from different cellular sources can activate endothelial cells (17, 18), a major cellular player in blood vessel angiogenesis and plaque formation (19, 20), we queried whether EVs derived from monocyte/platelet aggregates could impact on HUVEC reactivity. An overnight protocol was applied, testing initially a concentration-range of 1 to 20 EVs per endothelial cell. These experiments combined with published data (17, 21) indicated that a ratio of 10 EVs/Cell was optimal for our experimental approach. Indeed, microscopy imaging showed changes in HUVEC morphology after incubation with EVs isolated from monocyte stimulated with TNF-α (Fig. 3a). Furthermore cells showed a significant uptake of these EVs after 24 hour incubation (Fig. 3b). Next we quantified markers of cell activation.

**Figure 3.**
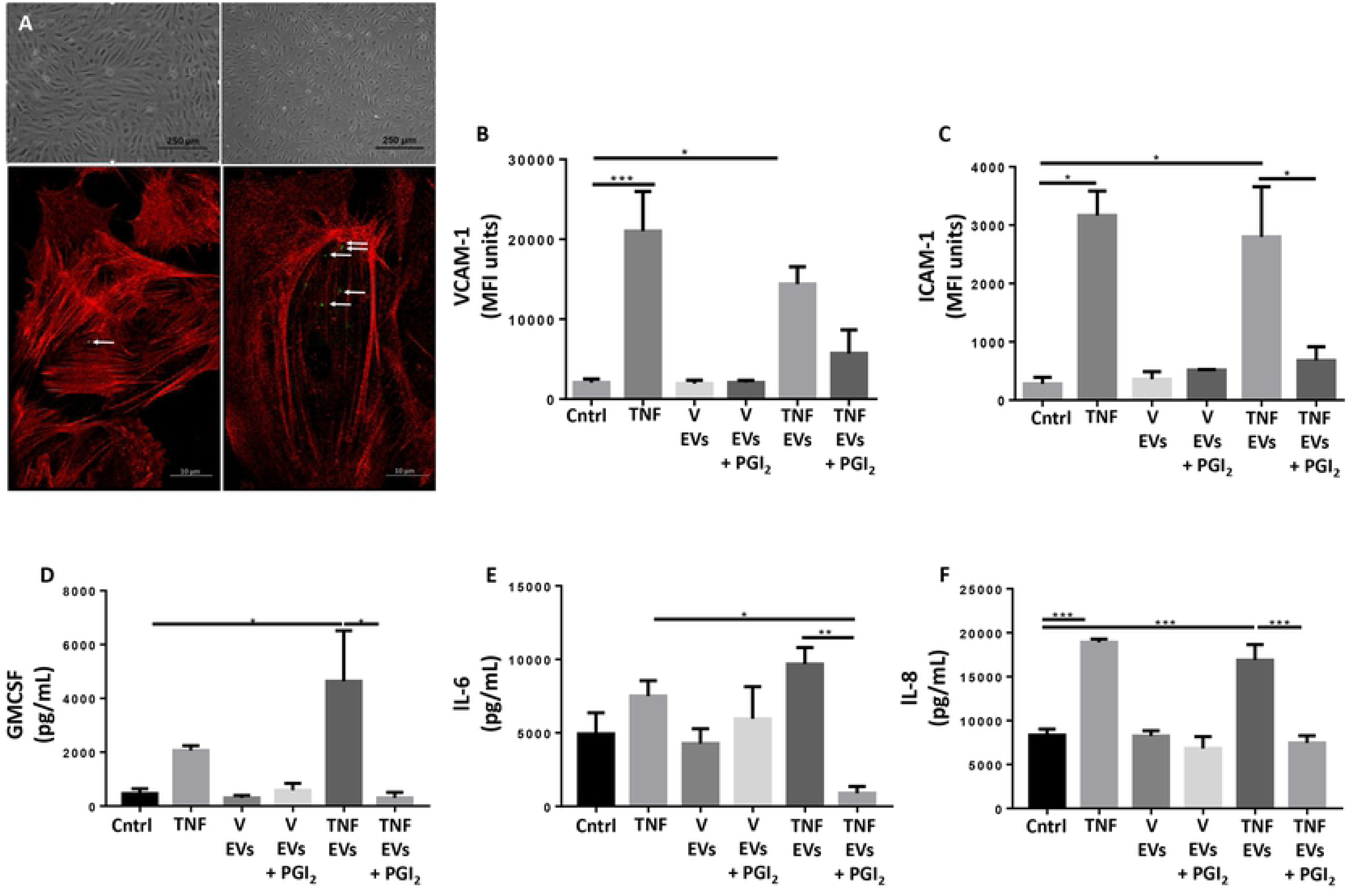
Monocyte/platelet EVs activate HUVEC *in vitro*. Monocyte were obtained as in Figure 2 and incubated with vehicle (V) or TNF-α (50 ng/ml), in presence or absence of Iloprost (1 µM; PGI_2_) for 60 min. HUVEC were incubated with the reported EVs (10×10^6^/ml) overnight. Cells were stained for flow cytomentry analysis and and supernatants were collected and analysed for cytokine release. (a) Representative microscopic image of HUVEC after treatment monocyte EVs from vehicle-incubated monocyte or TNF-α stimulated cells. (b) Confocal images of the uptake by HUVEC stained with Phalloidin (red) after 24 hours of BODIPY labelled EV isolated from TNF-α stimulated cells (green/white arrows). (c) Quantification of ICAM-1 and VCAM-1 expression (MFI units) in HUVEC treated with different subsets of monocyte derived EVs. (d) Quantification of GMCSF, IL-6 and IL-8 levels by ELISA. (*p<0.05, **p<0.01, ***p< 0.001; one-way ANOVA post Bonferroni test, mean ± SEM of n=5 cell preparation incubated with distinct EV preparations from different donor cells).

Flow Cytometry analysis revealed that expression of ICAM-1 and VCAM-1 was significantly upregulated when cells were treated with 10 ng/mL of TNF-α as positive control. Furthermore, similar increases were recorded when HUVEC were stimulated with EVs isolated from monocytes enriched preparations incubated with TNF-α. Of interest, ICAM-1 levels were no modified at all following incubation with EVs isolated from Iloprost and TNF-α stimulated monocytes (Fig. 3c). When similar experiments were repeated with the same concentrations (10×10^6^) of platelet EVs, isolated from cells in both stimulated (TNF-α) or in resting conditions, expression of either ICAM-1 or VCAM-1 was not modified, suggesting a synergistic role of monocyte/platelet aggregates in releasing functional EVs upon TNF-α stimulation (Fig. S4). Of relevance, only negligible amounts of residual TNF-α were detected in any of the vesicle preparations used (Fig. S5).

Cytokine measurements of HUVEC supernatants was then conducted. Cell incubation with EVs released by monocytes stimulated with TNF-α augmented concentrations of GM-CSF, IL-6 and IL-8 to a significant degree (Fig. 3d-f). When EVs were produced in presence of Iloprost, a lower regulation of these three cytokines was quantified (Fig. 3-f). These data together suggested a different pro-inflammatory effect of EVs generated from enriched monocyte preparations in response to several conditions chosen to mimic vascular inflammation.

### EV triggers differential activation of human atherosclerotic plaque

Having confirmed that EVs derived from monocyte/platelet aggregates can activate endothelial cells, we tested if they might be a functional determinant in atherosclerosis. Thus, we assessed their function on an atherosclerotic plaque using an *ex-vivo* organ culture protocol (Fig. 4a-b). Herein we compared an overnight incubation with EVs generated from different cellular activation protocols, using the same concentration of EVs, as described in the previous section, to resemble vascular inflammation. Then, we quantified cytokines and proteins released in the supernatants from the plaque fragments.

**Figure 4.**
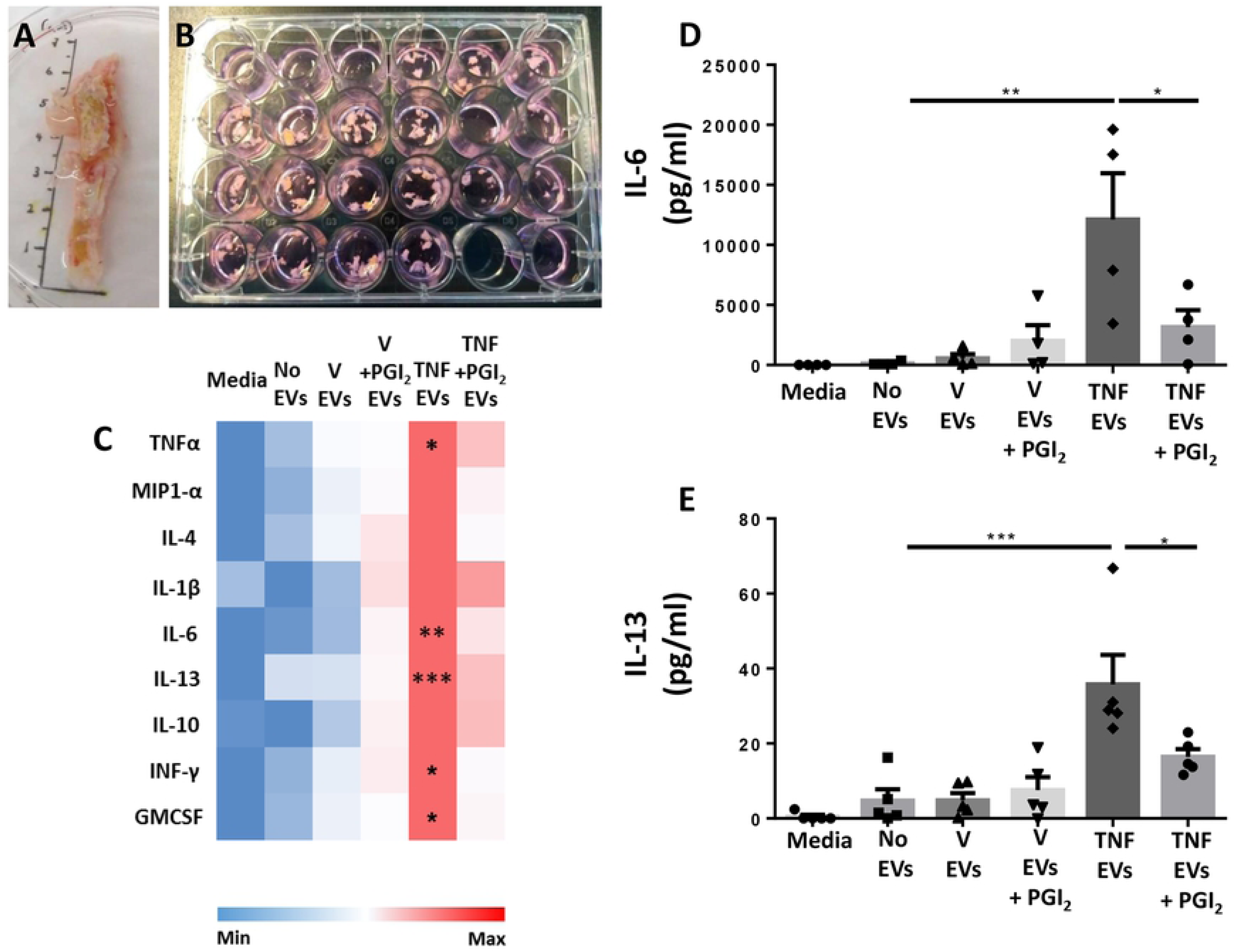
Monocyte/platelet EVs activate human atherosclerotic plaque ex-vivo. Monocyte were obtained as in Figure 2 and incubated with vehicle (V) or TNF-α (50 ng/ml), in presence or absence of Iloprost (1 µM; PGI2) for 60 min. Human atherosclerotic plaque fragments were incubated with the reported EVs (10×106/ml) overnight. Supernatants were collected and used for ELISA analysis. (a,b) Representative images of human femoral plaque and fragment incubation. (c) Heat map analysis showing qualitative modulation of cytokine release (linear scale bar). (d,e) Quantification of IL-6 and IL-13 levels by ELISA. (*p<0.05, **p<0.01, ***p< 0.001; one-way ANOVA post Bonferroni test, mean ± SEM of n=5 plaques incubated with distinct EV preparations from different donor cells).

Cytokine multiplex analyses revealed that treatment of the plaque with EVs released by monocytes stimulated with TNF-α, augmented concentrations of TNF-α, IL-6, IL-13, IFN-γ and GM-CSF in the culture media (Fig. 4c and Supplementary Table S2). As mentioned already above only negligible amounts of residual TNF-α were detected in any of the vesicle preparations used (Fig. S5). When EVs were generated in the presence of Iloprost, a much milder regulation of the general cytokine response was noted (Fig. 4c). These findings seemed to confirm the acquisition of a pro-inflammatory phenotype of EVs not only *in vitro* but also *ex vivo* when monocytes enriched preparation were stimulated with TNF-α. Such an effect was markedly attenuated when EVs were generated by Iloprost+TNF-α treatment, a finding corroborated by further quantification of IL-6 and IL-13 in the supernatants. (Fig. 4d,e). Of importance, the use of 0.1% FBS to enable plaques fragments viability did not affect the experimental outcome.

To acquire a view of the broader effects of these EVs on plaque reactivity, tissue conditioned media were analysed by proteomics. In total, 654 proteins were identified in human plaque supernatants as reported in Table S3. Subsequently, we performed statistical analysis in order to determine proteins differentially secreted between plaques treated with different EVs subsets as compared to the untreated plaque. In total, 52 proteins resulted significantly modulated with the majority of these proteins (13) being uniquely modulated when the plaque was treated with TNF-α-stimulated monocyte/platelet aggregates EVs. Treatment of plaque with EVs from unstimulated cells (vehicle) produced a limited response with 6 significant proteins being modulated. Both EVs from Iloprost (PGI_2_) and Iloprost+TNF-α monocytes resulted in the modulation of the release of a different set of 8 unique proteins (Fig. 5a). Only 3 proteins were common between all the groups (Fig. 5a): guanine nucleotide-binding protein subunit beta-2-like 1 (GNBL1; a regulator of several signalling pathways), Serpin B6 (SERPINB6; natural inhibitor of serine proteinases) and Voltage-dependent anion-selective channel protein 1 (VDAC1; a protein that forms a channel through plasma membrane). Details of the proteins identified in this analysis are reported in Table S4. As shown in Figure 5a, functional enrichment analysis of the modulated proteins mapped to distinct cellular functions including immune system, neutrophil degranulation and extracellular matrix organization. Of interest, plaque fragments treatment with EVs isolated from TNF-α stimulated monocytes not only increased the overall number of modulated proteins but a higher number of proteins involved in these functions emerged as well.

**Figure 5.**
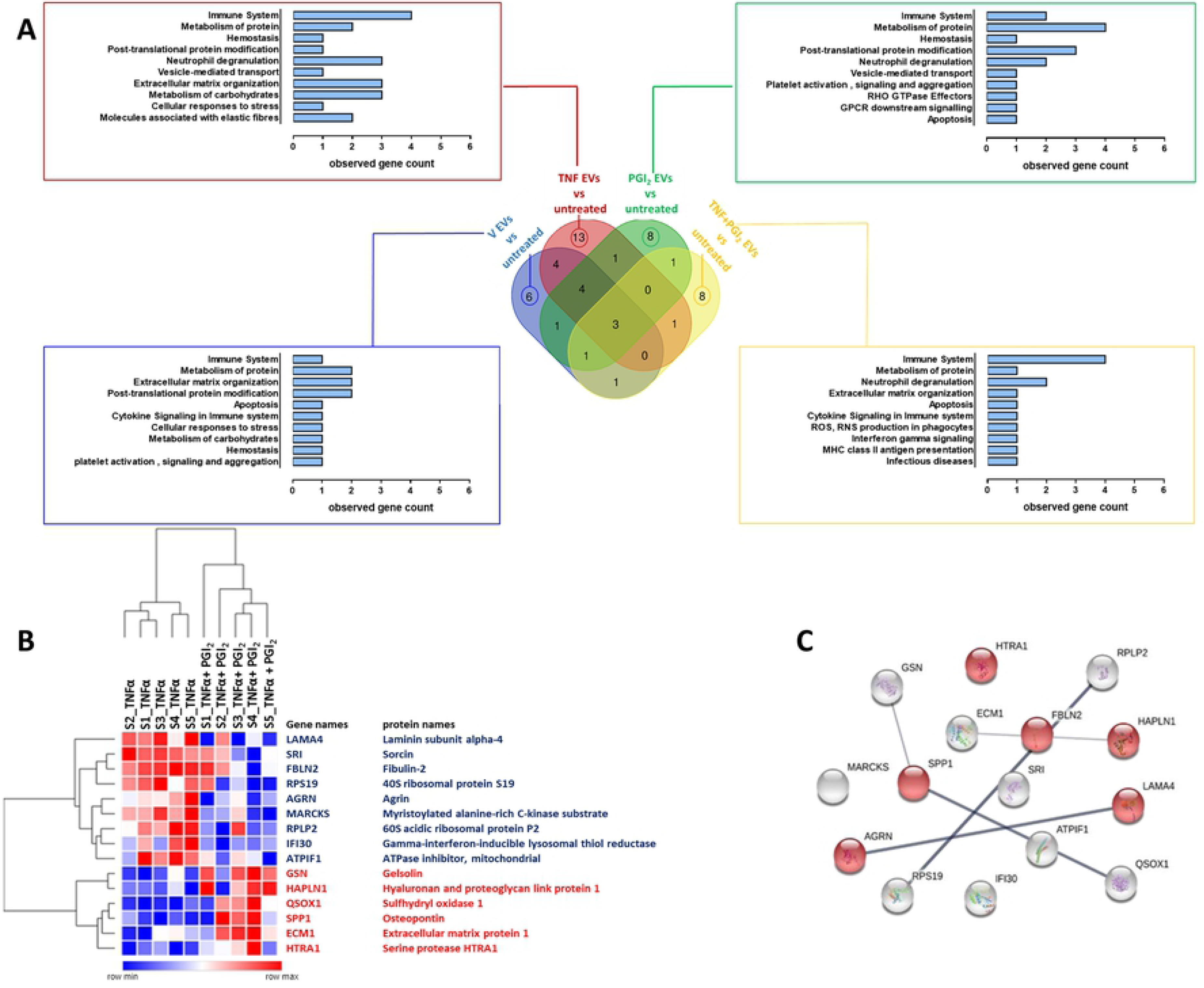
Modulation of secreted proteins from human atherosclerotic plaques by monocyte/platelet EVs. Monocyte were obtained as in Figure 2 and incubated with vehicle (V) or TNF-α (50 ng/ml), in presence or absence of Iloprost (1 µM; PGI_2_) for 60 min. Human atherosclerotic plaque fragments were incubated with the indicated EV subsets (10×10^6^/ml) overnight and proteomic conducted on supernatants. (a) Venn Diagram and related reactome pathway enrichment analysis obtained by PANTHER software clustering the significantly modulated proteins (p<0.05) by the EV treatments as indicated. (b) Hierarchical clustering heatmap identifying the 15 secreted proteins that were significantly modulated when comparing TNF-α EVs versus TNF-α+PGI_2_ EVs (p<0.05). Red represents up-regulated proteins while blue depicts down-regulated proteins. (c) Protein-Protein interaction network of the 15 proteins obtained by STRING: network nodes represent proteins; network edges indicate the strength of data support; proteins associated with “extracellular matrix organization” pathway are highlighted in red.

Prompted by the distinct phenotypic response of the plaque to different EVs subsets, we performed a targeted analysis directly comparing plaque treated with TNF-α EVs and Iloprost+TNF-α EVs: this analysis revealed a set of 15 modulated proteins as reported in Figure 5b. Of these, nine proteins were downregulated while six proteins were upregulated (Fig. 5b).STRING network analysis showed an enrichment in 6 proteins involved in extracellular matrix reorganization (Fig. 5c), a process which is crucial in vascular remodelling and atherosclerotic plaque formation. Fibulin (FBLN2) was consistently upregulated from plaques treated with TNF-α EVs but not with Iloprost+TNF-α EVs. This protein is emerging as a major effector in cardiac fibrosis and tissue remodelling (see Discussion) suggesting it may play a pivotal role in plaque activation and likely destabilization following incubation with TNF-α EVs. Conversely, Gelsolin (GSN) is a protein involved in actin filament assembly and organization (22), hence described to maintain the cytoskeleton structure in arteries (see Discussion), showed an opposite modulation.

### Characterization of monocyte EV subsets revealed differential protein expression associated with regulation of vascular inflammation and plaque formation

The experimental data presented so far are indicative of different pharmacodynamics properties produced by EVs obtained with TNF-α-treated monocytes compared to vesicles generated following treatment with Iloprost+TNF-α. In order to verify if these effects were mediated by a differential EVs composition we performed a proteomic characterization of TNF-α and Iloprost+TNF-α EVs. We identified 681 proteins in EVs by LS-MS/MS (Table S5), of which 32 proteins were significantly altered (p<0.05) when comparing TNF-α EVs to Iloprost+TNF-α EVs. (Fig. 6a): of these, 19 proteins were upregulated and 13 downregulated following cell incubation with Iloprost (Fig. 6b). Moreover, proteins uniquely expressed were also identified: 10 proteins for TNF-α EVs and only two for Iloprost+TNF-α EVs (Fig. 6a). Of interest, we detected Annexin A1, which is a faithful marker for membrane-spawn vesicles (23). Gelsolin (GSN) was identified as an interesting protein which was augmented in Iloprost+TNF-α EVs; this protein was also identified in the plaque proteomic analysis (see previous section). However, Fibulin was not a hit identified by this analysis (Table S5), while being regulated in the plaque proteome as discussed above. Next, and to further validate these data, we confirmed the relative abundance of a selected group of proteins by Western blotting and Imaging flow cytometry.

**Figure 6.**
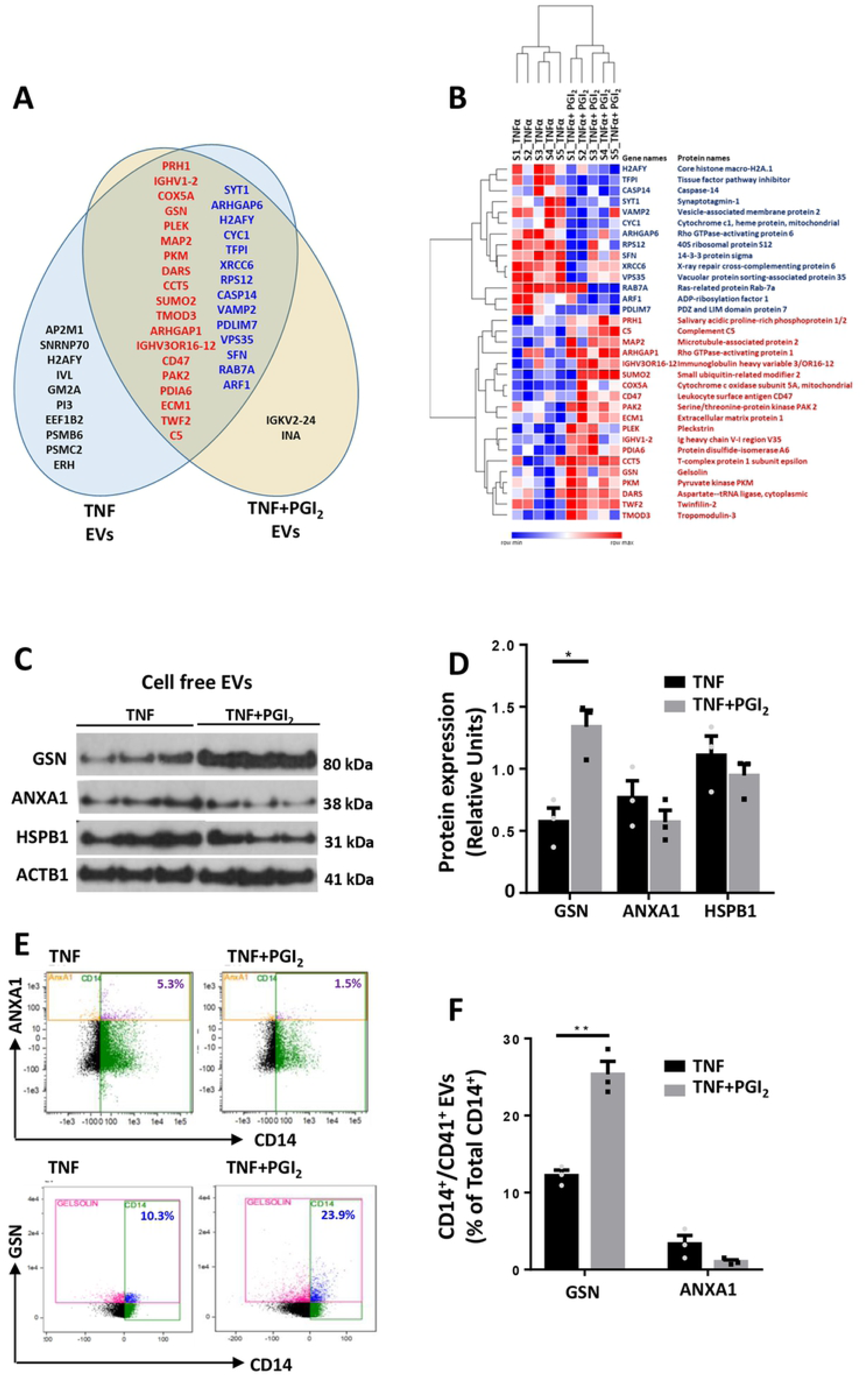
Proteomic analysis of monocyte/platelet EVs. Monocyte were obtained as in Figure 2 and incubated with TNF-α (50 ng/ml), in presence or absence of Iloprost (1 µM; PGI_2_) for 60 min, prior to EV purification. Targeted analysis highlighting differences between TNF-α and TNF-α+PGI_2_ EVs identified 33 proteins that were significantly altered (Table S5). (a) Venn diagram showing the proteins that are differentially expressed between TNF-α versus PGI_2_+TNF-α EVs. In the intersection of the diagram are reported the proteins there are significantly altered between the two EVs populations (p<0.05, red represents up-regulated proteins while blue depicts down-regulated proteins in TNF-α+ PGI2 EVs). In black, we identify proteins that are uniquely expressed in either EVs population. (b) Hierarchical clustering heatmap of differentially expressed proteins (centre of Venn diagram in panel a) between TNF-α versus PGI_2_+TNFα EVs (p<0.05). (c) Western blot analyses of distinct EV preparations used to detect immunoreactivity for Gelsolin (GSN), Annexin A1 (AnxA1), heat shock protein β1 (HSPB1) and β-actin (ACTB1). Three distinct EV preparations were tested. (d) Densitometry analysis, ACTB1 was used as loading control. (e) ImageStream™ analysis of a select group of proteins identified in the proteomic screen (see Methods for details). (f) Expression levels of GSN and ANXA1 in CD14+ EVs. *p<0.05, **p<0.01, ***p< 0.001; Mann Whitney test, mean ± SEM, n=3 distinct EV preparations.

To this end, equal numbers of monocyte/platelet EVs of each subset were loaded and immunostained for GSN, ANXA1, HSPB1 employing ATCB as a loading control. The blots confirmed that GSN was enriched in Iloprost+TNF-α EVs (Fig. 6c), whereas HSPB1 and ANXA1 were mildly regulated across the two EV subsets (Fig. 6d), again confirming the proteomics results. ImageStream analyses revealed that GSN and ANXA1 were also detected of the surface of the EVs (Fig. 6e), establishing again the selective enrichment of GNS in EVs isolated from monocytes stimulated with Iloprost and TNF-α (Fig. 6f), but not major changes for ANXA1. When CD41+ EVs were analysed, GSN+ EVs were 46.1±1.94% in TNF-α EVs and 63.4 ± 2.15% in the Iloprost+TNF-α EV group (mean ± SEM, n=3 distinct preparations).

To determine the cell source of these exemplar proteins, surface staining and intracellular staining of human monocytes and platelets aggregates was performed by microscopy. While ANXA1 was selectively expressed, to a high abundance, by monocytes (Fig. 7a,b), the majority of GSN seemed to be expressed by platelets both intracellularly and on their cell surface (Fig. 7a,b), while only a small amount was associated to monocytes likely because adherent to platelets. Similar results were also shown by Western blot when the same proteins were investigated in platelet or monocyte (the latter containing residual platelets) lysates. Loading of decreasing concentrations of monocyte and platelet whole lysates revealed GSN to be highly expressed in platelets, and as described by the immunofluorescence results, only a small amount of it was detected in the monocyte lysates possibly because of platelet contamination (Fig 7c). Conversely, monocyte lysates contained a consistent higher amount of ANXA1 represented by both the 38kDa and the 34kDa bands (Fig 7c). Interestingly, platelet extracts displays only minimal amounts of the cleaved form of ANXA1 (Fig 7d).

**Figure 7.**
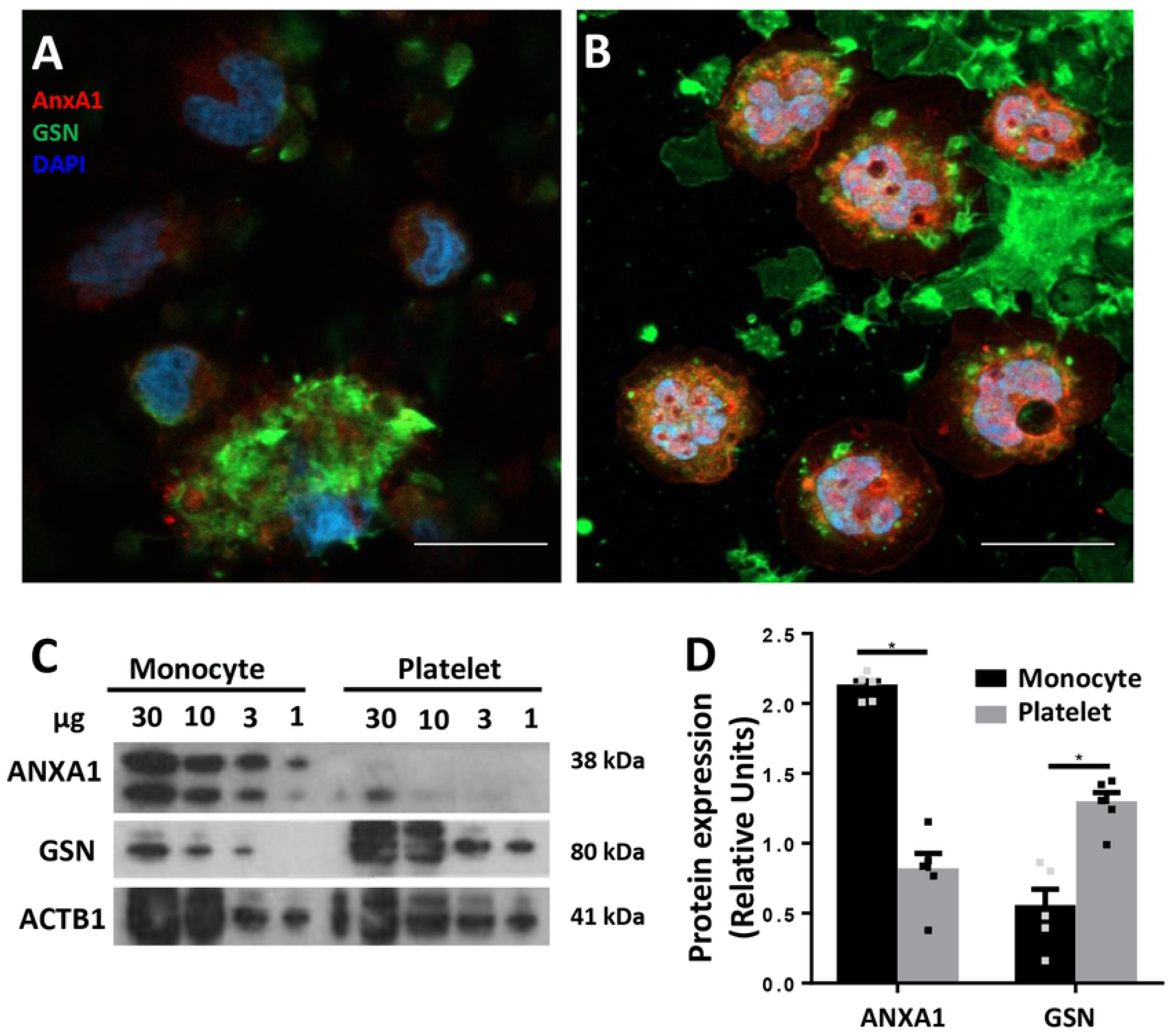
Selective expression of Annexin A1 (ANXA1) and Gelsolin (GSN) in human monocytes and platelets. The monocyte preparation, that contains platelet, was prepared as in Figure 2. Cells were permeabilised or left intact prior to staining for ANXA1 and GSN, prior to imaging. (a) Surface staining for ANXA1 (red) and GSN (green) in monocytes/ platelets aggregates. (b) Intracellular staining for ANXA1 (red) and GSN (green) in monocytes/ platelets aggregates. DAPI counterstaining (blue) indicates cell nuclei. Representative of three distinct cell preparations. Scale bar = 10 μm. (c) Western blot analysis for ANXA1 and GSN in monocyte and platelet lysates. (d) Densitometry analysis, ACTB1 was used as loading control. (*p<0.05, **p<0.01, ***p< 0.001, Mann Whitney test, mean ± SEM of n=5 with different donor cells).

In supplementary Table S6 we compare our data with other previously published proteomics data conducted on EVs from THP-1 cells (a surrogate of monocytes) and platelets.

## Discussion

In this study we provide evidence that monocyte/platelet-derived EVs are by and large pro-inflammatory and can activate not only endothelial cells, but also the atherosclerotic plaque, inferring a pathogenic role in *in vivo* settings. We further identify some subtlety in relation to the mode of activation of the monocyte with particular attention to the presence of an aggregated and/or adherent platelet. Using a pharmacological approach to preferentially attenuate platelet reactivity, we could produce EVs with a lower pathogenic impact, at least in the context of an atherosclerotic plaque activation. These different outcomes were not related to physicochemical features of the EVs but rather to their composition as indicated by the proteomic analysis. Since transient aggregates between monocytes and platelets can form in several settings of vascular inflammation, we propose that this inter-cellular cross-talk can generate EVs which may extend the patho-physiological relevance of this event. Clinical management with anti-platelet therapies may have beneficial effects also through a modulation of the quality of EVs released from monocytes.

Monocyte/platelet aggregates are a reported feature of vascular inflammation, being identified in several pathological settings, both in man and experimental animals. As an example, an experimental medicine study following kidney transplantation, revealed that addition of a 4-week anti-platelet therapy to immunosuppressive drugs reduced monocyte/platelet aggregates as well as several other markers of vascular inflammation (24). These aggregates have also been reported in stroke (25) and in heart failure (26). In heart failure, a specific subset of monocyte/platelet aggregates was negatively correlated with better prognosis, indicating a direct or indirect role for the aggregates in promoting sustained damage, or reduced repair, of the cardiac tissue. Finally, circulating monocyte/platelet aggregates have been detected in hypertension, where an independent predictor for their formation was systemic blood pressure (27), and in coronary artery disease. In the latter condition, monocyte/platelet aggregates increase in patients compared to healthy controls (28, 29), an increase quantified to be more than two-fold (30). In all these studies, the pro-atherogenic properties of the aggregates has been suggested. Of interest, whereas some of the studies summarised here proposed an initiating role of the platelets (24, 30), there is evidence in the context of coronary artery disease that indicates monocyte activation as the inciting event, leading to the formation of these aggregates (31).

In all cases, monocyte/platelet aggregates and more generally leukocyte/platelet aggregates, are transient in their association and dissociation. As a relevant example, Furman *et al* demonstrated that numbers of circulating monocyte/platelet aggregates in patients with acute myocardial dysfunction were higher within the first 4 hours of acute coronary symptoms and gradually returned to basal values in the 4-8 hour period post-infarct (32). In a longer prospective study about patients undergoing elective coronary bypass surgery, the numbers and reactivity of monocyte/platelet aggregates decreased to basal level after 3 months from surgery (33). Thus we reasoned that EVs could be represent a viable way to monitor longer-term effects of aggregates formation, either as a biomarker or as *bona fide* effectors of pathogenesis. As such we took advantage of the presence of platelets in the preparations of monocytes purified from human whole blood. A subtle and sophisticated role for the platelet emerged in these experimental conditions, whereby attenuation of platelet activation with the prostacyclin analogue did not affect platelets adhesion to the monocyte, while reducing the generation of EVs. In parallel experiments, we used PAF. This stimulus activated preferentially the platelet, produced a much larger number of EVs and specifically of CD41+ EVs: all these effects were significantly inhibited by prostacyclin. This set of results validated our conclusion that Iloprost acted predominantly on the platelet, yet it was able to affect EVs when a myeloid cell stimulus like TNF-α was applied. Monitoring cytokines released from the monocytes unveiled the inhibitory effect of the Iloprost which added further substance to this conclusion. Altogether we have a system where with TNF-α, we stimulate the monocyte predominantly, but there is a ‘co-stimulatory’ action attained by the adherent platelets. These data are in agreement with a proposed cross-talk in plaque formation and progression, whereby the platelet adherent to the monocyte favours migration of the leukocyte into the plaque, which would then develop to macrophages (34–36). Platelet-delivery of cholesterol could feed forward the process of macrophage differentiation into a foam cells (37). Here we reasoned that one downstream result of platelet/monocyte aggregate formation would be production of pro-inflammatory EVs.

When EVs generated from the different *in vitro* incubation protocols were analysed by Nanosight™ a relatively similar size was measured, with no major differences in median and mode of the distribution when TNF-α or PAF were applied as stimuli, in the presence or absence of prostacyclin analogue. In all cases, an average diameter of >100 nm was quantified suggesting that EVs produced are predominantly formed by membrane-spawn vesicles and not by exosomes (23). This evidence was corroborated by the proteomic analysis and further validated by Western blot, which identified ANXA1 in all subsets of EVs, with no significant changes across the groups. A recent elegant study identified ANXA1 as a genuine marker for membrane-spawn vesicles, also referred to as microparticles (23). More interesting to us is the emerging evidence that the same cell can generate EVs which are at least in part different in relation to the stimulus applied or the microenvironment. Our own work on neutrophil-derived EVs reported major functional differences between EVs produced in suspension versus in adhesion settings, with the former being reparative and anti-inflammatory (Dalli et al. 2013; Headland et al. 2015) while the latter EVs are mainly pro-inflammatory (6, 39). In line with this, in this study, we could demonstrate that monocyte-derived EVs isolated from mixed platelet/monocyte aggregates in inflammatory conditions (TNF-α) bind to and are internalized by HUVECs. Importantly, EVs isolated from TNF-α treated monocyte/platelet aggregates, but not from untreated cells or from cells previously treated with Iloprost, up-regulated ICAM-1 and VCAM-1 expression and increased the release of GM-CSF, IL-8 and IL-6 from endothelial cell monolayers. Before exploring further the differences in composition we further tested the potential different effector functions of TNF-α and Iloprost+TNF-α EVs using an organ culture protocol. For all the reasoning summarised above we focused on the atherosclerotic plaque.

There has been quite some interest in EVs and atherosclerosis, mainly with a focus on vesicles released from the plaque, possibly as a downstream determinant of pathogenic processes operative within the plaque. Several studies showed that EVs are mainly derived from leukocytes and i) are endowed with thrombogenic activities (40), ii) can increase intra-plaque neovascularization and plaque vulnerability, mainly iii) because they enhance proliferation of endothelial cells and angiogenesis (41) through the presence of tissue factor activity (42). The leukocyte origin was further confirmed by Mayr *et al*, using a combination of unbiased analyses to quantify and qualify the myeloid EV fraction, as well as EVs from smooth muscle cells and erythrocytes. Metabolomics performed in the same study showed an increase in taurine, further emphasizing the monocyte/neutrophil-produced oxidative microenvironment within the atherosclerotic plaque (14). Here we revealed marked modulatory functions of monocyte EVs applied to the plaque. After 24 h incubation, the plaque was relatively viable as assessed with both focused and unbiased approaches, the former being multiple cytokine quantifications, the latter proteomic analysis, both conducted on plaque supernatants. The induction of several cytokines by TNF-α EVs is indicative of strong pro-inflammatory actions supporting the hypothesis that if generated within the plaque (perhaps from the extravasating monocyte bearing platelets on its surface) or migrated to the plaque, these vesicles can fuel local inflammatory processes. The increase in IL-6 is remarkable and fits with several studies that identify the importance of this cytokine in the development of atherosclerosis. Exogenously administered IL-6 significantly enhances (∼5-fold) the development of fatty lesions in mice (43). The pathogenic properties of IL-6 through enhancing endothelial dysfunction and aortic stiffness was demonstrated in rheumatoid arthritis patients treated with the IL-6 receptor inhibitor tocilizumab: the neutralizing anti-IL-6 therapy successfully reduced articular inflammation and decreased endothelial dysfunction, measured as impaired flow mediated dilatation and aortic stiffness by pulse wave velocity (44). A recent Mendelian randomized study, focusing on the single nucleotide polymorphisms in the IL-6 receptor gene, highlighted loss of function as a viable approach for the prevention of coronary heart disease (45). It was of great interest to us that the vesicles generated by the monocyte preparation stimulated with Iloprost+TNF-α displayed a totally different impact on the plaque. The cytokine response of the plaque was essentially blunted when compared to that quantified following overnight incubation with TNF-α EVs.

The fact that TNF-α EVs markedly affected the reactivity of ex vivo cultured atherosclerotic plaque was further confirmed by proteomic analysis of the conditioned media with the identification of a 52 significantly modulated proteins in response to the different EVs subsets. In particular, the proteome of the plaque conditioned media unveiled modulation of a different group of proteins between TNF-α and Iloprost+TNF-α EVs with an important distinction. While TNF-α EVs augmented ∼3-fold the levels of fibulin-2 in the plaque supernatants, Iloprost+TNF-α EVs failed to do so, indicating a head-to-head difference in line with the cytokine analyses. Fibulin-2 is an extracellular matrix protein that has been positively associated with fibrosis of the myocardium (46). In a model of heart failure, mice nullified for fibulin-2 are less susceptible to fibrosis while developing hypertrophy to the same extent as their wild type counterparts. Such an outcome is secondary to transforming growth factor beta expression in the presence of fibulin-2, together with potentiation of its signalling in target cells. Moreover, while selective expression of this protein in the aortic arch vessels it is known already to be associated with the morphogenic events that regulate heart development, in post-natal life fibulin-2 is also produced by endothelial cells of the coronary arteries and veins (47). These properties dovetail with the results obtained by our experiments whereby modulation of fibulin-2 follows the inflammatory status of the plaque and the regulation afforded by distinct types of EVs. Furthermore, the fact that fibulin-2 is required for higher deposition of collagen-I and collagen-III in cardiac fibrosis corroborate the pathogenic role that this protein may have in plaque formation and/or progression. One would derive that strategies to modulate fibulin-2 levels within the plaque, perhaps through delivery of antisense or blocking strategies with natural or semi-synthetic vesicles (47, 48), could be a viable therapeutic approach to impact on the progression of atherosclerosis.

Finally, the experiments presented and discussed above justified an in-depth analysis of the potential differences between TNF-α EVs and Iloprost+TNF-α EVs. While we recognise that structural lipids, lipid mediator precursors (49), microRNA and other nucleic acids (50) could vary between the two vesicle types, as a proof-of-concept for fundamental differences in composition, we analysed their protein contents. In general a lower number of significantly modulated proteins were detected in Iloprost+TNF-α EVs compared to TNF-α EVs suggesting that attenuation of platelet activation not only reduced the number of CD14+ and CD14/CD41+ EVs, but also modified the actual composition of these microstructures. Addition of Iloprost reduced the number of proteins exclusively identified in monocyte EVs from 10 proteins to 2. Comparison with published proteomic lists revealed interesting overlaps. As an example, THP-1 monocytic cells stimulated with lipopolysaccharide yield EVs that contain EEF1B2 (an elongation factor) and PSMC2 (proteasome subunit) (51), two of the proteins uniquely identified here for TNFα EVs. Out of six studies of platelet EVs, we focused on two studies (52, 53) were similar preparation protocols were applied for the generation of the EVs. Thus, INA (cytoskeleton component) uniquely identified in Iloprost+TNF-α EVs was identified by Pienimaeki-Roemer *et al.* in senescent platelet EVs (52). Platelet EVs also express PSMC2 as well as AP2M1 (vesicle transporter) and PSMB6 (another proteasome subunit) (52). Similar overlaps were noted for the proteins modulated in both subgroups of EVs analysed here, as reported in Table S6. An interesting hit was gelsolin, detected in more abundance in Iloprost+TNF-α EVs and also in plaques treated with this EV subset: this protein is endowed with anti-inflammatory properties and has been identified in resolving inflammatory exudates (54). In this study, addition of gelsolin to chondrocytes exerted positive modulation of extracellular matrix protein deposition while inhibiting metalloproteases and other catabolic enzymes. In the context of an atherosclerotic plaque, such a profile would yield a stabilizing effect. In fact, published data using proteomic approaches have reported gelsolin downregulation in atherosclerotic coronary arteries compared to pre-atherosclerotic coronaries and mammaries (55). The same authors demonstrated how reduction in gelsolin levels caused i) cytoskeleton deregulation within the human atherosclerotic coronary media layer and ii) switch of medial vascular smooth muscle cells from a contractile to a synthetic phenotype (proinflammatory) (56). Furthermore, circulating levels of gelsolin are reduced in patients with a diagnosis of asymptomatic carotid artery plaque (57) and in patient with ankylosing spondylitis undergoing TNF-α antagonist-infliximab therapy when compared to healthy matched controls (58).

Collectively these data strengthen the close relationship between monocytes and platelets for the generation of EVs which may be of mixed origin, with proteins that may derive from one cell or the other. As such future studies may focus on the biogenesis of EVs from the aggregates and perhaps reveal a common budding process into the vesicles that emerge from them.

In conclusion, the activating effect of monocyte-derived vesicles on the reactivity of the atherosclerotic plaque reflects the contribution of platelet adhesion. Monocyte/platelet aggregates, accepted as a predictive marker of several cardiovascular pathologies including coronary artery disease, may have longer lasting pathogenic effects through generation of vesicles which may propagate pro-inflammatory actions. Modulation of platelet reactivity could help attenuating the detrimental properties of these vesicles.

## Material and methods

### Monocyte purification and flow cytometry characterization

All volunteers gave written, informed consent to blood collection and the procedure was approved by the Queen Mary Ethics of Research Committee (QMERC2014.61) for healthy controls. Blood (30 ml) was drawn from healthy volunteers using a 19G butterfly needle with tourniquet applied and anticoagulated with 0.32% w/v sodium citrate. To inhibit platelet activation, Iloprost™ (2µM; stable prostacyclin analogue; Sigma-Aldrich, Gillingham, UK) was added to whole blood prior cell separation. The blood was, then centrifuged at 150 x*g* for 20 min and the platelet rich plasma (PRP) removed and replaced with PBS+1mM EDTA. Following another centrifugation step, RosetteSep™ cocktail (15028, StemCell Technology, Vancouver, Canada) was added (50 µl/ml of blood) and samples rested at room temperature for 20 mins. Blood was then diluted 1:1 with PBS+1mM EDTA and layered over 15 ml Histopaque 1077 (Sigma-Aldrich, Gillingham, UK), centrifuged for 20 min at 1200 x*g* room temperature to separate monocytes from other cells. The monocyte layer was harvested and washed at 300 x*g* for 10 min. Following another washing step, the monocyte pellet was re-suspended in phenol red-free RPMI (Gibco, Waltham, US) and the concentration adjusted as needed.

For peripheral blood mononuclear cells (PBMCs) isolation, whole blood was centrifuged at 130* ×g* for 20 minutes and plasma was removed. For every 30 ml of whole blood, erythrocytes were depleted by sequentially layering 10 ml PBS followed by 8 ml of 6% w/v dextran (high molecular weight, Sigma-Aldrich, in PBS) and gently inverting. After 15 min, the leukocyte-rich fraction was collected and layered over Histopaque 1077 and centrifuged for 30 minutes 450 x*g* at room temperature to separate granulocytes from PBMC. PBMCs were washed once by centrifuging at 300 *×g* and resuspended in RPMI for further use. For polymorphonuclear cells (PMN) isolation, whole blood was centrifuged at 130 x*g* for 20 min, plasma removed, erythrocytes depleted on 6% w/v dextran (high molecular weight, Sigma-Aldrich, in PBS). Then, the leukocyte-rich fraction was layered over Histopaque and centrifuged for 30 min 450 x*g* at room temperature. The PMN layer was harvested, washed and cell concentration adjusted as needed.

After isolation cells were treated with Fc receptor blocking solution and stained with anti-CD14-APC (2 μg/ml, 61D3; Biolegend, San Diego, USA), anti-CD41-PE (2 μg/ml, HIP8; Biolegend), anti P-selectin-FITC or anti-PSGL1-PE (2.5 and 1.5 μg/ml, AC1.2 and KLP-1 respectively; Becton Dikinson, Franklin Lakes, USA). Cells were acquired on an LSR Fortessa cytometer.

For platelet isolation, PRP was isolated directly from blood by centrifugation as above. PRP was further processed into washed platelets (WP) by addition of 2μg/ml Iloprost and 0.02U/ml apyrase (M0398S, NEB), prior to centrifugation at 1000 *xg* for 10 minutes. Pellets were re-suspended in modified HEPES Buffer containing Iloprost and apyrase and washed a second time. Platelet where then counted, and concentration was adjusted to 3×10^8^/mL before stimulating.

### Fluorescent microscopy analysis of monocytes and platelets

Isolated monocytes containing platelets were spotted on Alcian blue-coated glass slides and fixed in cold 4% paraformaldehyde (4 °C, 30 min). After fixation, cells were washed with PBS and then blocked in PBS with 0.2% BSA (for surface staining) or PBS with 0.1% Triton and 0.2% BSA (T-PBS; for intracellular staining) for 30 min at room temperature shaking. Following blocking, monocytes and platelets were incubated with primary specific antibodies against Annexin A1 (ANXA1; 5 μg/ml; clone 1B, in house generated) and Gelsolin (GSN; 1.54 μg/ml clone EPR1942; Abcam, Cambridge, UK) in either PBS+0.2% BSA or T-PBS+0.2% BSA overnight at 4 °C. The cells were then washed and incubated with secondary antibody Alexa Fluor 488 anti-rabbit (5 μg/ml, Molecular Probes Invitrogen, Eugene, USA) or Alexa Fluor 592 anti-mouse (5 μg/ml, Molecular Probes Invitrogen) in T-PBS+0.2% BSA for 1 h at 20 °C shaking. Cells were then mounted with a glass coverslip using Fluoroshield™ Histology Mounting Medium with DAPI (Sigma-Aldrich) and visualized under the microscope Zeiss LSM800 Imaging System.

### Western blot analysis of monocytes and platelets

Presence GSN and ANXA1 was confirmed through standard SDS-PAGE (Millipore, Watford, UK), loading extracts from 30µg, 10 µg, 3µg and 1 µg of isolated monocyte or washed platelet lysates. Western blot was conducted with specific antibodies against ANXA1 (ANXA1; 5 ng/ml; clone 1B), GSN (1.54 ng/ml clone EPR1942; Abcam, Cambridge, UK), or anti-β-actin (ACTB; 5 ng/ml; clone AC-74, Sigma-Aldrich) overnight at 4 °C followed by a 1 h incubation with either an HRP-conjugated goat anti-mouse IgG or goat anti-rabbit IgG (Dako, Cambridge, UK). Proteins were detected using Luminata™ Forte Western HRP Substrate (Millipore, Watford, UK) visualized on Hyperfilm™ (GE Healthcare, Buckinghamshire, UK).

### Generation and isolation of monocyte EVs

Monocytes (1×10^6^ cells/mL) were incubated with TNF-α (50 ng/mL; Sigma-Aldrich), platelet activator factor (PAF, 1 μM; Cayman Chemical, Ann Arbor, USA) or PBS for 60 min at 37°C. Washed platelets were incubated with 50 ng/mL TNF-α for 20 min at 37°C. When the prostacyclin analogue Iloprost was added, it was used at 1 µM (Iloprost®; Sigma-Aldrich). Cell suspensions were centrifuged at 4,400 x*g* at 4°C for 15 min to pellet cells and/or platelets, followed by a second centrifugation at 13,000 x*g* at 4°C for 2 min to remove remaining contaminants (e.g. apoptotic bodies). EVs were enriched by centrifuging at 20,000 x*g* at 4°C for 30 min, the supernatant was removed, and pellets were re-suspended in filtered sterile PBS.

### Characterization of monocyte-derived EVs

#### Nanoparticle tracking analysis for sizing EVs

Approximately 0.5 ml of EVs (between 10^6^ to 10^8^ vesicles) in suspension were loaded onto the Nanosight NS300 with 488 nm scatter laser and high sensitivity camera (Malvern Instruments Ltd., Malvern, UK); five videos of 90 seconds each were recorded for each sample. Data analysis was performed with NTA2.1 software (Nanosight, Malvern, UK). Software settings for analysis were the following, Detection Threshold: 5–10; Blur: auto; Minimum expected particle size: 20 nm.

#### ImageStream™ analysis for quantification and characterisation of EVs

EVs were analysed and counted using fluorescence triggering on an ImageStreamx™ MKII imaging cytometer as described previously (Headland et al. 2015). Briefly, vesicles were labelled with 50 μM BODIPY maleimide fluorescein or BODIPY Texas-Red (Life Technologies, Carlsbad, USA), and acquired as such or after labelling with either 2 μg/ml anti-CD14-APC (61D3; Biolegend), 2 μg/ml anti-CD41-PE (HIP8; Biolegend) or one of the following Pacific Blue or Alexa Fluor 488 conjugated antibodies: anti-Annexin A1 (ANXA1; 1 μg/ml; clone 1B), anti-Gelsolin (GSN; 0.1 μg/ml clone EPR1942; Abcam). Fluorescence minus one (FMO) controls were used for gating all protein antigen-positive events. Approximately 20,000 events were acquired per sample.

#### Proteomic analysis of EVs

EVs derived from monocytes treated with TNFα in presence or absence of Iloprost® were pelleted at 20,000 *xg* for 30 min, resuspended in 20 μl ice cold RIPA buffer containing protease inhibitor (Sigma Aldrich). Protein content was measured by spectrophotometry (Nanodrop 2000, ThermoFisher Scientific, Waltham, USA) selecting Protein A280 program and 50 µg of proteins were used for trypsin digestion. Mass spectrometry analysis of the proteins obtained from EVs was performed on tryptic digests obtained using the Filter Aided Sample Preparation protocol as previously described (59). EVs proteome profile was determined by LC-MS/MS analysis as previously described in the methods section All data and materials have been made publicly at the PRIDE (60) Archive (EMBL-EBI) with the dataset identifier PXD014325.

#### Western blotting analyses

Presence of a select group of proteins identified by proteomic analysis was confirmed through standard SDS-PAGE, loading extracts from ∼30 × 10^6^ EVs per lane (Millipore, Watford, UK). Western blot was conducted with specific antibodies against ANXA1 (ANXA1; 5 μg/ml), GSN (1.54 μg/ml clone EPR1942; Abcam), anti-Heat shock protein β-1 (HSPB1; 5 μg/ml; clone G3.1; Abcam), or anti-β-actin (ACTB; 5 μg/ml; clone AC-74, Sigma-Aldrich) overnight at 4 °C followed by a 1 h incubation with either an HRP-conjugated goat anti-mouse IgG or goat anti-rabbit IgG (Dako). Proteins were detected using Luminata™ Forte Western HRP Substrate (Millipore) visualized on Hyperfilm™ (GE Healthcare).

### Experiments with Human Umbilical Vein Endothelial Cells (HUVEC)

#### Isolation and culturing of HUVEC

Cells were freshly isolated from umbilical cords that were kindly donated by the midwifery staff of the Maternity Unit, Royal London Hospital (London, UK) with an approved protocol (East London & The City Local Research Ethics Committee reference 05/Q0603/34 ELCHA). Cells were cultured 0.5% gelatin coated T75 flasks, in 5% CO_2_ at 37°C with complete Medium 199 (Gibco, Waltham, USA) containing 100 U penicillin, 100 mg/mL streptomycin, and 2.5 µg/mL fungizone (Gibco) supplemented with 20% Human serum (Sigma-Aldrich, UK), and used up to passage 4.

#### Assessment of adhesion molecule expression by flow cytometry

HUVEC were grown to confluence in 6 well plate coated with 0.5% gelatin and stimulated 24 hours with TNF-α (10 ng/mL), or 10×10^6^ EVs isolated from monocyte-platelet aggregates subsequent to stimulation with vehicle, TNF-α, or iloprost+TNF-α, in 0.5% human serum complete media, as described above. After isolation cells were treated with Fc receptor blocking solution and stained with anti-ICAM-1-PE (1 μg/ml, HA58; Biolegend, San Diego, USA), anti-VCAM-1-BV711 (0.5 μg/ml, 5110C9; Optibuilt, USA). Cells were acquired on an LSR Fortessa cytometer.

#### Confocal imaging of EV uptake

HUVEC were seeded overnight on 0.5% Gelatin coated µ-Slide 8 Well Glass Bottom (80826, Ibidi) at a concentration of 1×10^5^. Cells were stimulated with 1×10^6^ EV previously stained with 2.5 µM BODIPY-FITC for 20 min and pelleted at 20,000 *xg* for 30 minutes at 4°C. Subsequently, the cells were fixed with 4% PFA for 15 min and blocked and permeabilised with PBS containing 2% BSA and 0.1% Triton-X for one hour. Cells were finally stained with 1.5 nM Phalloidin AF647 (A22287, ThermoFisher Scientific) for 45 minutes. A Nanoimager-S microscope (ONI, UK) was used for microscopy of the HUVEC using ONI software. The following excitation/emission conditions were used in conjunction with x100 magnification oil immersion objectives: BODIPY 488/561 and AF647 640/658. The images acquired were analysed using supplied ONI and ImageJ software packages. Distinct BODIPY fluorescent (green) points identified by eye in the micrographs were considered as distinct EV.

### Experiments with the human atherosclerotic plaque

#### Isolation and ex vivo culture

Patients with clinical and angiographic evidence of atherosclerosis undergoing revascularization surgery were recruited to the study. All 5 patients undergoing carotid or femoral endarterectomy gave written informed consent. The study was approved by the Ethics Committee of St. Vincent’s University Hospital in Dublin, and in accordance with the International guidelines and Helsinki Declaration principles. Surgical atherosclerotic plaque samples were harvested in physiological saline. After dissection, they were stimulated in 24-well plates for 24 hours at 37°C, 5% CO_2_ in RPMI with 0.1% exosome depleted Fetal Bovine Serum with the different EV subsets (10×10^6^ per well), which were isolated from monocyte-platelet aggregates subsequent to stimulation with vehicle, TNF-α, or iloprost+TNF-α as described above. After 24 hr incubation, tissues samples and supernatants were collected and snap frozen in liquid nitrogen for subsequent analysis by mass spectrometry analysis and multiplex ELISA assay.

#### Proteomic analysis of plaque supernatants

Conditioned media obtained from plaque supernatants (from plaques treated with or without EVs, n=5) was defrosted at room temperature and centrifuged firstly at 14,000 x*g* for 2 min to remove debris and then at 20,000 x*g* at 4°C to remove residual EVs from both the stimulation and the exosome depleted FBS, this sequential centrifugation reduced further ∼94% the number of EV contained in the FBS. An equal volume of 20% trichloroacetic acid was added to the sample and incubated on ice for 1 hr. Samples were centrifuged at 10,000 x*g* for 15 min at 4°C and washed with 500 μl of ice-cold acetone. After 5 min incubation, proteins were spun for 5 min at 5,000 x*g* at 4°C, acetone was removed and pellets were left to dry. Dried protein pellets were re-suspended in 8 M Urea/ 25 mM Tris– HCl, pH 8.2. Disulphide bonds were reduced with 5 mM Dithiothreitol (DTT) and protected with 15 mM iodoacetamide. Proteins were digested with sequencing grade trypsin (1:100; Promega, USA) overnight at 37°C and peptides concentration was checked by spectrophotometry (Nanodrop 2000; ThermoFisher Scientific, Waltham, USA). Then,15 μg of peptides were purified using ZipTipC18 pipette tips according to manufacturer’s instructions (Millipore, Billerica, USA), resuspended in 2% Acetonitrile/0.1% formic acid solution, prior to injection of 2 µg of purified peptides into an Ultimate3000 nano-LC system coupled to a Q Exactive mass spectrometer (ThermoFisher Scientific) for mass spectrometry. Peptides were separated by increasing acetonitrile, 2 to 33%, in a linear gradient of 40 min on a C18 reverse phase chromatography column packed with 2.4 µm particle size, 300 Å pore size C18 material (Dr Maisch GmbH, Ammerbuch Entringen, Germany) to a length of 120 mm in a column with a 75 µm, using a flow rate of 250 nL/min. All data were acquired with the mass spectrometer operating in an automatic data dependent acquisition mode (DDA, shotgun). A full mass spectrometry service scan at a resolution of 70,000, AGC target 3e6 and a range of m/z 350–1600 was followed by up to 12 subsequent MS/MS scan with a resolution of 17,500, AGC target 2e4, isolation window m/z 1.6 and a first fix mass of m/z 100. Dynamic exclusion was set to 40 s. Mass spectrometry data were processed using label-free quantitation method in MaxQuant software v.1.3.0.547, using the human Uniprot database (release 2016_3). All data and materials have been made publicly available at the PRIDE Archive (EMBL-EBI)(60) partner repository with the dataset identifier PXD014324.

Downstream analysis of proteomic data was performed by Perseus software (version 1.6.0.7). Only the proteins present in at least 50% of the samples in at least one group (“Untreated” plaque, “TNF-α”, “PGI_2_”, “TNF-α+PGI_2_ EVs treated plaque) were considered identified. Proteins found to be differentially expressed between groups (Student’s T-test P<0.05, FDR 0.05) were subjected to enrichment analysis and were distributed into categories according to cellular component, molecular function, biological process, KEEG pathways and reactome pathways using PANTHER (Version 14.1) or STRING Database (Version 10.5). STRING was also used to generate protein-protein interaction networks.

#### Multiplex ELISA analysis of plaque and monocyte supernatants

GM-CSF, IL-6 and IL-8 concentrations from HUVEC conditioned media or GM-CSF, IFN-γ, IL-1β, IL-4, IL-6, IL-10, IL-13, MIP-1α and TNF-α concentrations from the centrifuged conditioned media of the human plaques were measured by enzyme immunoassay using commercially available human 96 well-plate multiplex kit for tissue culture samples (MSD, Gaithersburg, USA) according to the manufacturers’ guidelines. The same cytokines plus MCP-1 instead of TNFα were quantified in the monocyte conditioned media following removal of EVs by centrifugation.

### Statistical analysis

All statistical analysis and graphing were performed in GraphPad Prism 6 Software, IDEAS 6.2 for Image Stream Plots and FlowJo V6 for LSRFortessas Plots. Data are expressed as mean ± standard error (SEM) unless stated differently. Analyses applied to the different experimental data are indicated in each figure legend. A p value of < 0.05 was considered significant to reject the null hypothesis.

### Data Availability

Mass spectrometry proteomics analysis of monocyte-derived EVs: PRIDE PXD014324 (http://www.ebi.ac.uk/pride/archive/projects/PXD014324) Mass spectrometry proteomic analysis of human plaque supernatant: PRIDE PXD014325 (http://www.ebi.ac.uk/pride/archive/projects/PXD014325).

## Acknowledgments

We thank Mr Joseph Dowdall and Mr Stephen Sheehan, Department of Vascular Surgery, St Vincent’s University Hospital for the provision of material and we thank Jan Nagenborg and Antonino Cacace for logistic support. The authors acknowledge the support of the UCD Conway Institute Core Technology mass spectrometry facilities.

EVOluTION has received funding from the European Union’s Horizon 2020 research and innovation programme under the Marie Sklodowska-Curie grant agreement No. 675111 (S.O., M.P.). Wellcome Trust (programme 086867/Z/08/Z) to M.P. The ImageStream™ used was funded by the Wellcome Trust (infrastructure grant 101604/Z/13/Z). M.deG. is supported by an IRC Government of Ireland postdoctoral fellowship (IRC GOIPD/2017/1060), E.P.B and C.G are supported by Science Foundation Ireland grants 15/US/B3130 and 15/IA/3152 and a strategic research award from JDRF NY, USA. This work has been facilitated by the National Institute for Health Research Biomedical Research Centre at Barts Hospital NHS Trust.

## Author contributions

M.P. devised the project, the main conceptual idea and proof outline. S.O. planned the project, designed and performed experiments, analysed the data. C.G., helped supervise the project, designed experiments, analysed the data; D.C. and L.V.N. designed experiments, analysed the data. M. deG. and E.P.B assisted with *ex vivo* atherosclerotic plaque experiments and S.M. helped carry out the proteomic determinations and analyses. T.M.M. helped with the interpretation of the proteomic data. M.B. helped providing the human samples used in the study. S.O. and M.P. wrote the manuscript.

## Conflict of interest

None.

